# The BLMP-1 transcription factor promotes oscillatory gene expression to achieve timely molting

**DOI:** 10.1101/2021.07.05.450828

**Authors:** Yannick P. Hauser, Milou W.M. Meeuse, Dimos Gaidatzis, Helge Großhans

## Abstract

Gene expression oscillators can coordinate developmental events in space and time. In *C. elegans*, a gene expression oscillator directs rhythmic accumulation of ∼25% of the transcriptome, and thus thousands of transcripts, presumably to control molting, a process of rhythmic skin regeneration. Recently, a reverse genetic screen identified several transcription factors important for molting. Here, we characterize one of these, BLMP-1, orthologous to the mammalian transcription repressor PRDM1. We find it to be important for timely molting, and oscillatory gene expression. We propose a dual function for BLMP-1 in shaping oscillatory gene expression and coupling it to a set of direct targets, which ensures cuticular integrity. With mammalian PRDM1/BLIMP1 promoting regular cycles of postnatal hair follicle regeneration, our findings point to the possible existence of a fundamentally conserved clock mechanism in control of rhythmic skin regeneration.

## Introduction

Biological ‘clocks’, based on genetic oscillators, drive dynamic processes ranging from circadian rhythms (Patke et al., 2020) to vertebrate somitogenesis (Oates et al., 2012). Yet, it is unclear whether similar processes operate in other instances of cyclical developmental processes. For instance, nematodes undergo a recurring process of skin barrier renewal, termed molt. Molting involves the synthesis of a new cuticle from the single-layer epidermis during a period of behavioral quiescence, termed lethargus, and the subsequent shedding of the old cuticle (ecdysis), which demarcates the end of a larval stage (Lazetic and Fay, 2017) (Fig 1A). In analogy to flies, where pulses of ecdysone trigger molts, it may be hypothesized that steroid hormone signaling controls the onset of the molt by triggering a gene expression cascade, and at least two transcription factors of the nuclear hormone receptor family, NHR-23 and NHR-25, function in molting (Lazetic and Fay, 2017). However, both the identity of the hypothetic hormonal signal and the mechanism that would specify its timely release, i.e., the underlying temporal control mechanism, have remained elusive. Indeed, although many factor have been identified that are required for successful molting (Lazetic and Fay, 2017), it is not known that any of these specifically affect the timing of molts.

**Fig 1:**
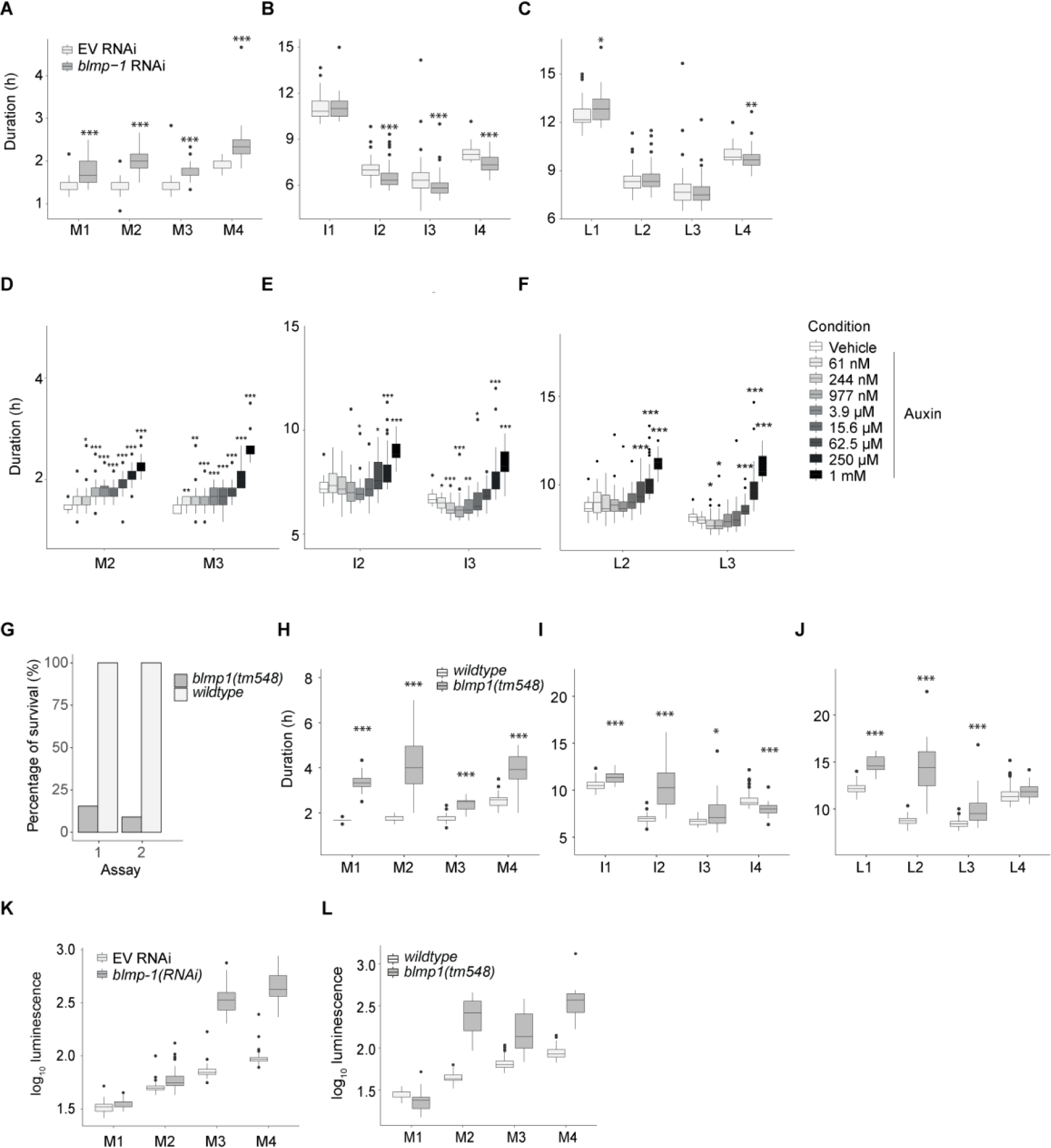
BLMP-1 loss of function leads to developmental defects. **A – C,** Boxplots of molt (A), intermolt (B) and larval stage (C) durations in mock (n=59) and *blmp-1* RNAi-treated animals (n=50) as determined by a single-animal luciferase assay. Significantly different durations are indicated (* P<0.05, ** P<0.01, *** P<0.001, Welch two-sample and two-sided t-test) **D – F,** Boxplots of molt (A), intermolt (B) and larval stage (C) durations of *aid::blmp-1* animals exposed to vehicle or auxin at the indicated concentration from egg stage to adulthood. **G,** Bar plot of manual quantification of lethality (inferred from loss of luminescence, Fig. S1) detected in two replicate luciferase assays for wild-type (replicate 1: n=76, replicate 2: n=80) and *tm548(tm548)* worms (replicate 1: n=129, replicate 2: n=77), respectively. **H – J** Boxplots of single animal molt (H), intermolt (I) and larval stage (J) durations of wild-type (n=76) and *blmp-1(tm548)* (n=20) mutant animals. *blmp-1(tm548)* animals are the surviving individuals from assay #1 in (G) (n=20). **K, L,** Boxplots showing log10-transformed luminescence intensities during the indicated molts in mock (EV) and *blmp-1* RNAi (K) and in wild-type and *blmp-1(tm548)* mutant animals (L). Significantly different durations are indicated in (A) – (F) * P<0.05, ** P<0.01, *** P<0.001, Wilcoxon unpaired, two-sample test. In the boxplots (A) – (F), (K) – (L), the horizontal line represents the median, hinges extend to first and third quartiles and the whiskers extend up to 1.5*IQR (interquartile range). Data beyond that limit is plotted as dots. See also Figure S1.

Consistent with the repetitive nature of *C. elegans* larval development, ∼25% of the larval transcriptome (corresponding to ∼3,700 genes, henceforth referred to as oscillating genes) exhibits high-amplitude oscillations (Grün et al., 2014; Hendriks et al., 2014; Kim et al., 2013; Meeuse et al., 2020) (reviewed in (Tsiairis and Großhans, 2021). The oscillating gene set is enriched for components of the cuticle, such as collagens, and factors involved in their processing or degradation, such as proteases. Their temporally choreographed expression may provide a basis for a robust and efficient execution of the molt (Tsiairis and Großhans, 2021). Moreover, as many other genes lack known functions in molting and/or exhibit expression peaks outside the molt, oscillations may represent a broader principle of temporal organization within larval stages.

Irrespective of peak phase, the expression of oscillating genes is generally synchronized with molting (Meeuse et al., 2020). This close coupling suggests a shared mechanism of temporal control. The fact that molting occurs with great regularity even in isolated animals, and without a need for obvious periodic environmental inputs implicates an intrinsic timing mechanism. At this point, it is unknown how this mechanism could emerge, but the precedence of circadian and somitogenesis clocks makes genetic oscillators, i.e., gene regulatory networks (GRNs) that generate oscillations through the wiring of their components, an attractive possibility. Yet, testing this model requires identification of potential oscillator components and an examination of their roles in oscillation period and developmental timing. Such components have remained largely elusive.

To identify candidate oscillator components, we recently screened through a set of 92 transcription factors with oscillating transcript levels to uncover those that caused abnormal numbers of timing of molts (Meeuse et al., 2022). The six hits thus identified included BLMP-1, whose depletion altered molt durations.

In *C. elegans*, BLMP-1 was previously shown to regulate gonad migration and terminal differentiation of epidermal cells (Horn et al., 2014; Huang et al., 2014). Recent work identified it as a pioneer factor involved in chromatin opening, which in turn might set the stage for the activity of rhythmically acting transcription factors (Stec et al., 2021). BLMP-1 is also required for an intact adult cuticle, likely through regulating the expression of particular collagen genes (Sandhu et al., 2021),

Here, we show that BLMP-1 loss causes extended larval molts and abnormal oscillatory gene expression. Functional BLMP-1 targets are mostly oscillating genes, including additional screen hits. Strikingly, titration of BLMP-1 degradation rates through auxin-induced degradation reveals that BLMP-1 depletion can accelerate molt onset, revealing a function in developmental timing, possibly through its effects on oscillatory gene expression. These findings provide a basis towards a mechanistic understanding of the *C. elegans* gene expression oscillator and its role in controlling developmental timing.

## Results

### Depletion of BLMP-1 alters developmental timing

Rhythmic transcription and RNAPII recruitment suggested an involvement of rhythmic transcription factor activity in generating transcript level oscillations. Recently, we exploited the coupling between mRNA oscillations and the molting cycle (Meeuse et al., 2020) and used a targeted RNAi-approach to identify transcription factors required for timely molting: screening through a set of 92 transcription factors with oscillating transcript levels we identified six that caused abnormal molting patterns (Meeuse et al., 2022). BLMP-1-depleted animals exhibited a phenotype of extended molt durations, which we decided to investigate in more detail here. We examined a larger number of animals using a luminescence-based assay that facilitates detection of molt entry and molt exit, and thus determination of molt, intermolt and larval stage duration, on single animals in high throughput (Meeuse et al., 2020; Olmedo et al., 2015). Briefly, animals expressing a luciferase transgene constitutively were cultured in a multi-well plate in a temperature-controlled luminometer in the presence of food and luciferin. Continuous light emission results, except during lethargus (molt), when animals do not feed so that luciferin uptake does not occur. Consistent with the screen results, *blmp-1(RNAi)* caused a lengthening of molts relative to mock-depleted animals (Fig 1A, Fig S1A,E).

Surprisingly, we observed in some but not all experiments largely unchanged larval stage durations, because lengthening of molts was counteracted by a shortening of intermolts (Fig 1A-C, Fig S1A-C, E-G). We wondered whether the variable effects on intermolt duration reflected differences in depletion achieved in different RNAi experiments. To achieve more controllable depletion of BLMP-1 than is possible by RNAi, we tagged the endogenous protein with an auxin-inducible degron (AID) in animals expressing *Arabidopsis* TIR1 ubiquitously (Zhang et al., 2015) to achieve a better control over degradation. Following addition of different amounts of auxin to embryos to trigger AID-BLMP-1 degradation, we indeed found an auxin dose-dependent effect: Increasing concentrations of auxins caused increasing molt durations, up to a maximum of a 2-fold increase at 1 mM auxin (Fig 1D). [These concentrations have little or no effect on animals lacking the degron on BLMP-1, (Meeuse et al., 2022)].

In contrast to molts, intermolts changed non-monotonically in response to different auxin doses: they decreased at lower auxin concentrations but increased at high auxin concentrations (Fig 1E). A largely constant, or under certain conditions and for some stages even shortened, larval stage duration resulted at lower auxin concentration, but an increased duration at higher auxin concentration (Fig 1F).

For *blmp-1(tm548)* mutant animals, considered null mutant (Horn et al., 2014; Huang et al., 2014), we frequently observed a gradual loss of luminescence after exit from the first molt (Fig S1M-O), which we interpret to reflect a developmental arrest or death (Fig 1G), although we have not investigated it further. For the subset of *blmp-1(tm548)* animals that developed through all larval stages, all four molts were greatly extended relative to wild-type animals (Fig 1H), whereas intermolts were more variably affected, with a substantial lengthening detectable for intermolt 2 (I2), but much less so for I1 and I3, and in fact a decrease for I4 (Fig 1I). From M1 onwards, molt, intermolt and larval stage durations also became more variable across individuals (Fig 1H-J).

Whereas luminescence is low in wild-type animals during lethargus, when luciferin is not ingested, we consistently observed elevated luminescence during M2 through M4 in both *blmp-1* RNAi-treated and *blmp-1(tm548)* mutant animals (Fig 1K,L, Fig S1D,H,L). Hence, a skin barrier defect, previously reported for adult *blmp-1* mutant animal (Sandhu et al., 2021), is also present in larvae (Fig S3P,Q). Collectively, these data reveal that BLMP-1 is required for normal temporal progression of development, while its absence compromises skin barrier function.

### BLMP-1 controls the expression of a large set of molting genes

To understand the molecular function of BLMP-1, we examined a publicly available BLMP-1 ChIP-seq dataset (GSE25803, modENCODE_DCC: 2612) to identify the genes that it binds. We identified 856 genes whose promoters (defined as a 1,000bp region centered around the ATG start codon) were bound by BLMP-1 in two independent experimental replicates (Fig S2).

Transcription factor binding tends to be poorly predictive of transcriptional regulation (Cusanovich et al., 2014). To define functional targets, we performed two replicate RNA sequencing time-courses on synchronized populations of *blmp-1(tm548)* mutant and wild-type animals grown at 25°C. To facilitate analysis and favor detection of direct consequences of BLMP-1 loss, we focused on early larval development, when developmental phenotypes are less pronounced (Fig 1, Fig S1), accepting that we may thereby underestimate the extent to which BLMP-1 affects the larval transcriptome. Moreover, to avoid false positives resulting from changes in developmental tempo (Fig S3A,B, (Tsiairis and Großhans, 2021)), we temporally realigned the gene expression of all time-courses.

Specifically, we plated synchronized early L1 stage larvae and sampled hourly over 24 h to cover the first intermolt, during which *blmp-1* mutant animals develop synchronously and at a largely normal rate, the greatly extended, yet still relatively synchronous molt 1, and an extended and variable intermolt 2, up to and into molt 2 (Fig 1E–G). Unlike in liquid medium in the luciferase assay, viability of *blmp-1* mutant animals appears largely unimpaired when grown on plate, a conditional lethality effect also observed by others (Stec et al., 2021).

Following alignment to the slowest time course, replicate 2 of the *blmp-1(tm548)* mutant time course (Fig S3, Methods), a dynamic threshold considering the magnitude as well as the significance of the effect (Fig. S4A, Methods) allowed us to identify among all expressed genes (n = 16,720) those that were differentially expressed between *blmp-1(tm548)* and wild-type (n = 1,655, Fig 2A).

**Fig 2:**
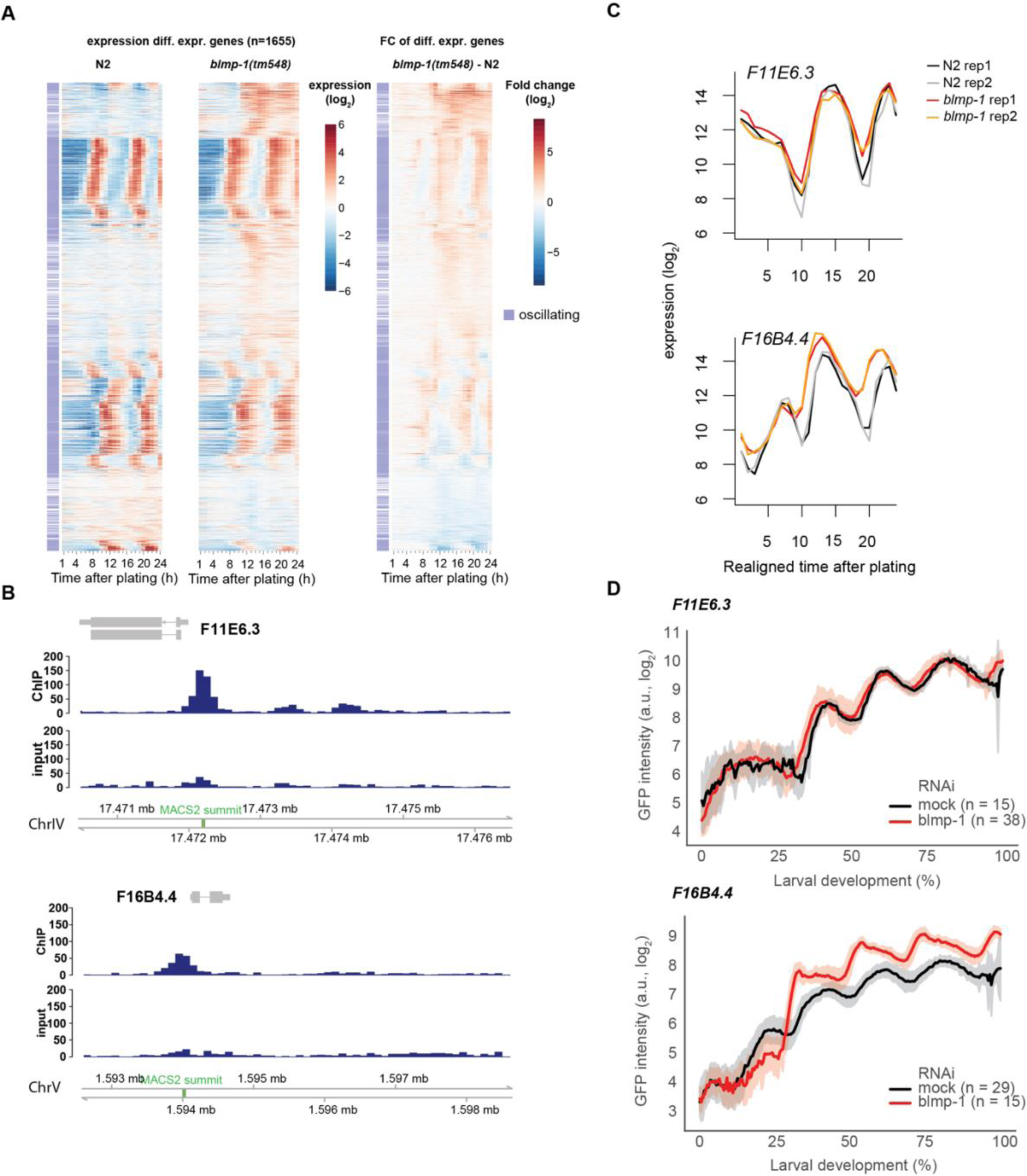
Time-course analysis reveals genes with dysregulated expression in *blmp-1(tm548)* animals. **A,** Heatmap of log_2_-transformed gene expression and fold changes of differentially expressed genes in realigned *blmp-1* mutant RNA sequencing time course. The heatmap was ordered according to a 1-dimensional tSNE (see Methods). **B,** BLMP-1 ChIP-seq binding profiles for the promoters of the genes shown in (C). The binding peak identified by MACS2 analysis is indicated in green. **C,** log_2_-transformed expression levels over realigned time for *F11E6.3* and *F16B4.4* transcripts for the indicated strains and replicates from RNA-sequencing time courses. **D,** Single worm GFP intensities of transcriptional reporters based on the promoters of the genes in (C) were scaled to larval development (hatch until molt 4 exit, see methods) and log_2_-transformed for mock (black) and *blmp-1* (red) RNAi-treated animals. The indicated number of single animals were followed over time; GFP intensities were averaged and plotted with the respective standard deviations in shading. See also Figures S3, S4

We confirmed the validity of the realignment approach through an orthogonal, single worm time-lapse imaging approach (Meeuse et al., 2020), which allowed us to observe larval stage durations, and thus developmental tempo, directly and in parallel to the expression of a given transcriptional *gfp* reporter. We selected the promoters of two genes, *F11E6.3* and *F16B4.4*, that were both bound by BLMP-1 (Fig 2B). However, consistent with the notion that transcription factor binding is not predictive of function, endogenous transcript levels of *F11E6.3* do not change significantly in the rescaled RNA-seq data (Fig 2C, Fig S3G), and this observation was replicated, and thus validated, by the transcriptional reporter: Expression dynamics were largely comparable for the *F11E6.3p::gfp* reporter in BLMP-1-depleted and mock control animals (Fig 2D, Fig S4B,C,F).

By contrast, BLMP-1 depletion affects both accumulation of the endogenous *F16B4.4* transcript and expression of an *F16B4.4p* reporter (Fig 2D, Fig S4D,E,G). Indeed, notably, both the shape and peak time of the *F16B4.4p* reporter expression changed substantially upon BLMP-1 depletion, validating the role of BLMP-1 in generating oscillatory gene expression patterns.

### Direct BLMP-1 targets are enriched for expression in tissues with strong blmp-1 expression

We intersected the BLMP-1 ChIP-seq and the RNA-seq data sets to define 248 genes as high-confidence BLMP-1 targets, based on BLMP-1 binding to their promoters and affecting their expression (Fig 3A). Base on published single cell expression data from wild-type animals (Cao et al., 2017), we found that these BLMP-1 target genes are preferentially expressed in tissues with high *blmp-1* levels, in particular seam cells, non-seam hypodermis, vulval precursor cells, excretory cells, socket cells and rectum (Fig 3B, Fig S5), distinguishing them from genes that were not significantly changed in the *blmp-1(tm548)* RNA sequencing time course (Fig 3C, Fig S5). We note that distal tip cells (DTCs) of the somatic gonad exhibited high *blmp-1* RNA levels but little or no evidence for an enrichment of the direct BLMP-1 targets identified by us, likely reflecting a limitation of bulk ChIP-seq and RNA-seq assays to detect such effects in a tissue comprising only 2 cells per animal.

**Fig 3:**
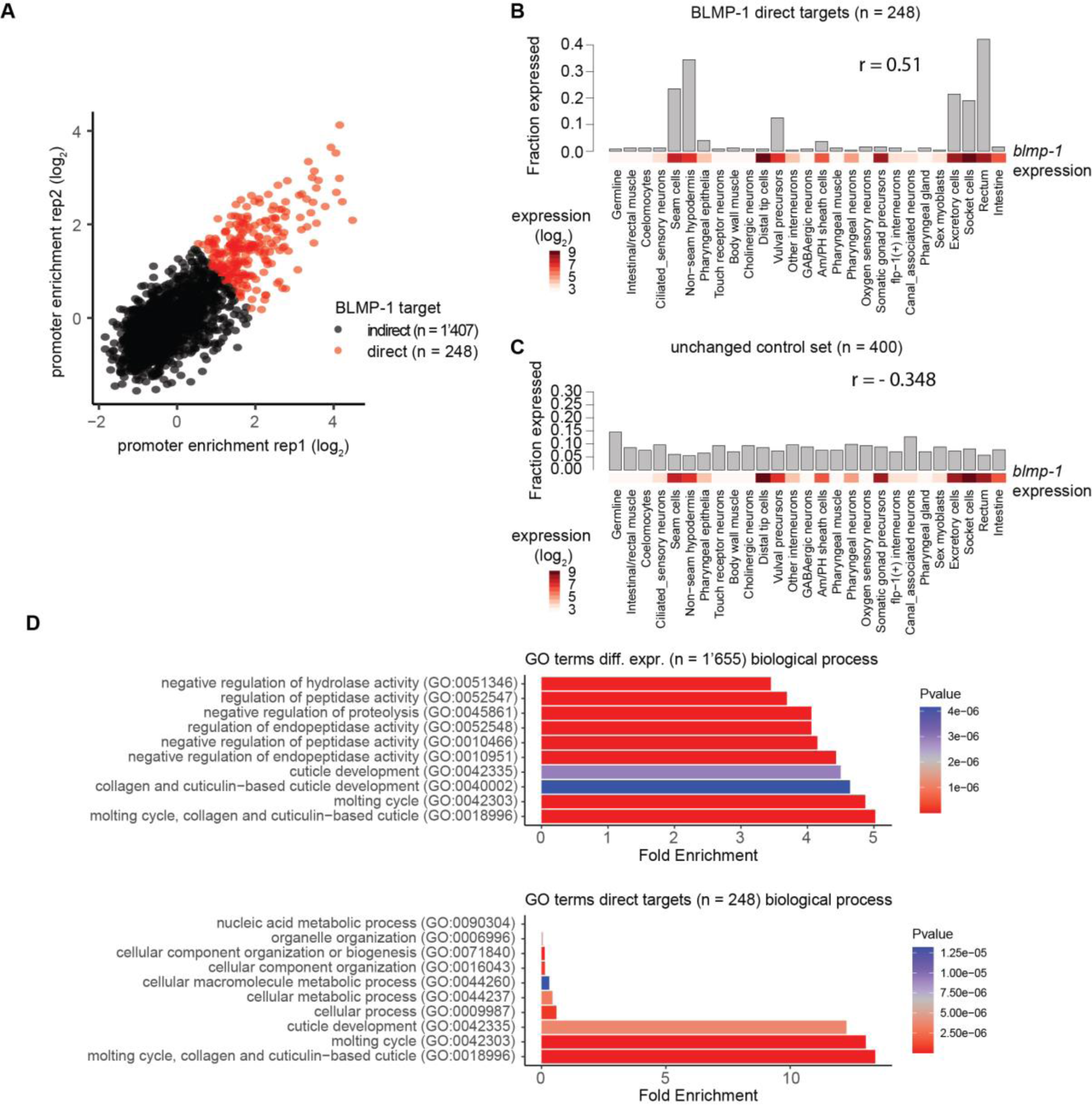
Direct BLMP-1 targets function in molting. **A,** Scatter plot showing promoter-binding of BLMP-1 as log_2_-fold enrichment over input in two replicate ChIP-Seq experiments for all genes that significantly change between *blmp-1(tm548)* and wild-type animals, shown in Fig 2A. Every dot is a gene, red indicates binding enrichment across replicates at a log_2_ average of >1. **B – C**, Fraction of direct target genes (B) and 400 non-significantly changed genes (selected by highest p-values) in the RNA sequencing time course (C) that are expressed at sufficiently high levels (log_2_ expression > 7) in individual cells (data from Cao et al., 2017). The Spearman correlation (r) of *blmp-1* expression with the fraction of genes significantly expressed in cells or tissues is indicated above the plot. **D,** Bar plots showing enrichment of gene ontology (GO) terms related to biological processes for genes differentially expressed in *blmp-1(tm548)* mutant animals (top, as in Fig. 5A) and direct BLMP-1 targets (bottom; as in panel (A)). The top 20 most highly enriched terms were sorted according to P-value followed by plotting the most 10 significant terms. See also Figure S5

We performed gene ontology (GO) analysis for differentially expressed genes, encompassing both direct targets and secondarily affected genes, as well as for direct targets only. We found that both sets exhibited a strong and significant enrichment of cuticle- and molting-related terms (Fig 3D), providing a possible molecular explanation of the molting defects that manifest in *blmp-1* mutant animals.

### BLMP-1 promotes oscillatory gene expression

We found a strong enrichment of oscillating genes in all BLMP-1-related categories relative to all unchanged genes, i.e., in BLMP-1-bound genes, in differentially expressed genes, and in direct BLMP-1 targets (Fig 4A,B). This effect was most pronounced for direct BLMP-1 targets, among which 220 out of 248 genes (88.7 %) were oscillating genes. These genes included *blmp-1* itself as well as two additional hits from the previous screen (Meeuse et al., 2022), *nhr-23* and *nhr-25*. Hence, regulation of oscillating gene expression is the predominant activity of this transcription factor.

**Fig 4:**
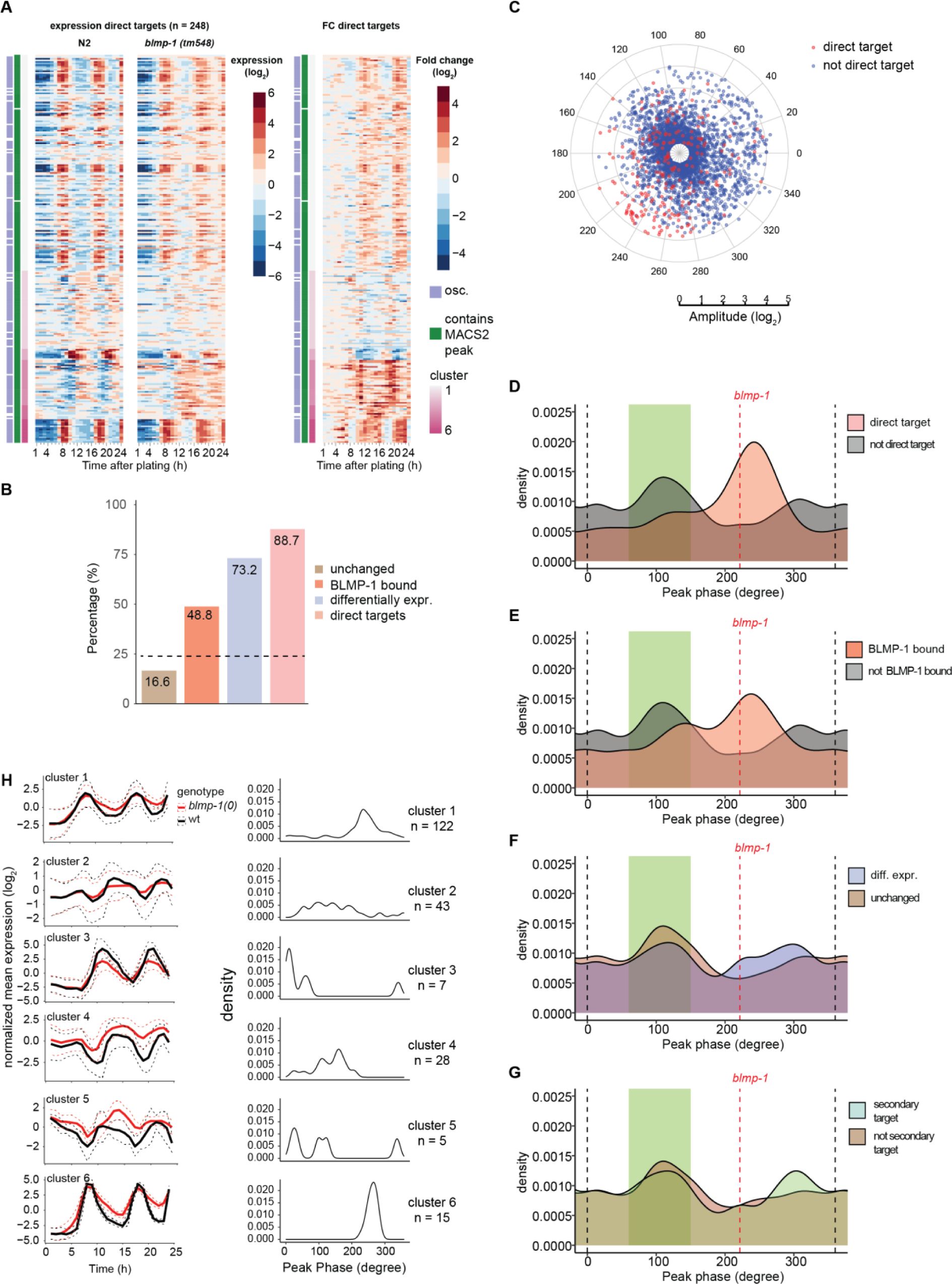
BLMP-1 targets shapes oscillating gene expression of direct targets. **A,** Heatmap of log_2_-transformed gene expression and fold changes of direct BLMP-1 targets as identified in Fig 3A. The heatmaps were all ordered according to hierarchical clustering based on the fold changes and cut into 6 different clusters, indicated in shades of purple. **B,** Percentage of oscillating genes among the indicated categories of genes. Categories are (right to left) “Direct targets” according to Fig 3A, “diff. expr.” (differentially expressed) according to Fig 2A, “BLMP-1-bound” according to Fig S2, and “unchanged”, which are all expressed genes used as input for differential expression analysis (see Methods) except those that are differentially expressed according to Fig 2A. Dashed line indicates the expected fraction of oscillating genes according to (Meeuse et al., 2020). **C,** Polar plot displaying the peak phase (circular axis, degrees) and the amplitude (radial axis, log_2_) of oscillating genes. The direct targets according to Fig 3A are marked in red. **D–G**, Peak phase distribution of oscillating genes among the indicated categories (as detailed in B). The phase of the molt is indicated in green. The dotted line at 0° and 360° indicate the boundaries of the circular phase information (360° = 0°). The peak phase of *blmp-1* mRNA is represented with the red dotted line. **H**, Mean-normalized mean expression and peak phase distributions of oscillating genes identified in the 6 clusters in A. The standard deviation is shown in dashed lines. See also Fig. S6

Direct, functional BLMP-1 targets had a strong preference for peak expression in interphase (Fig 4C, D) in wild-type animals. A similar, albeit smaller enrichment was also detectable for BLMP-1-bound genes that were not signicantly affect in gene expression by *blmp-1* mutation (Fig 4E). By contrast, when we looked at all differentially expressed genes (which includes both direct targets and secondary targets, which lack BLMP-1 promoter binding) or only at secondary target genes, this peak phase bias was greatly reduced (Fig 4F,G). Thus, BLMP-1 appears to have two, presumably connected, functions: It promotes oscillatory expression of its direct targets by modulating their transcription rhythmically. In addition, it promotes oscillatory expression of a larger set of genes that lack a clear peak phase signature and which it does not bind directly. These secondary targets thus appear to depend on promotion of core oscillator activity through BLMP-1. Consistent with this notion, oscillatory expression of secondary targets was greatly impaired in *blmp-1* mutant animals (Fig S6).

To explore mechanisms of direct target regulation, we used hierarchical clustering of the fold changes to resolve six groups of genes with distinct behaviors (Fig 4A,H). As for the secondary targets, a general impairment in oscillatory expression was evident, frequently manifesting in a reduction in oscillation amplitude in *blmp-1(tm548)* mutant animals (Fig 4H). Yet, the patterns differed for distinct clusters. Thus, in clusters 1 and 6, trough expression levels were up while peak levels remained unaffected. By contrast, in cluster 3 peak levels were decreased. (More complicated patterns in the other clusters may reflect a greater diversity in gene peak expression phases, diluting observable differences.) These findings indicate that BLMP-1 may act as both a transcriptional repressor and a transcriptional activator, depending on the identity of its target.

## Discussion

The biological functions and mechanism of the recently discovered *C. elegans* gene expression oscillations have thus far remained poorly defined (Hendriks et al., 2014; Kim et al., 2013; Meeuse et al., 2020). Here, building on a screen for candidate core oscillator components, we demonstrate that the BLMP-1 transcription factor promotes oscillatory gene expression and timely molting. BLMP-1 promotes rhythmic gene expression by both activating and repressing transcription of direct target genes, which include additional hits from the previous screen.

### BLMP-1 exhibits activating and repressing functions on direct targets to promote their oscillatory expression

Our analysis of *blmp-1* mutant animals revealed substantial transcriptional dysregulation preferentially affecting oscillating genes. Among the six clusters of direct BLMP-1 targets, i.e., genes that are dysregulated by *blmp-1* loss and normally bound by BLMP-1 in their promoters, we commonly observed reduced oscillation amplitudes. However, depending on the cluster, different effects contributed to this result: in some cases, peak expression levels decreased, whereas in others, trough levels increased. We emphasize that BLMP-1 binds to the promoters of all these genes, suggesting direct effects. Indeed, although Blimp1/PRDM1 proteins are well established transcriptional repressors (John and Garrett-Sinha, 2009), others recently reported a role of *C. elegans* BLMP-1 and mouse Blimp1 in transcriptional activation (Minnich et al., 2016; Stec et al., 2021). Our data support the notion that Blimp1/Prdm1 proteins may generally be capable of both repressing and activating target genes.

### BLMP-1 as a component of a C. elegans gene expression oscillator

Recently, (Stec et al., 2021) reported that BLMP-1 functioned as a pioneer transcription factor and suggested that it provide a permissive activity for dynamic gene expression. For instance, BLMP-1 binding would constitutively open chromatin at target loci to prime them for access by rhythmically active, instructive transcription factors. However, our quantitative analysis of the direct, functional targets of BLMP-1, i.e., the genes that are both bound and regulated by BLMP-1, revealed a clear peak phase preference, and thus a molecular signature of rhythmic BLMP-1 activity. Hence, our data support a model where rhythmic activity endows BLMP-1 with an instructive role in setting temporal gene expression and molting patterns. Incidentally, although a quantitative analysis does reveal a peak phase preference in BLMP-1-bound genes, this effect is indeed much more pronounced among functionally validated targets reaffirming the importance of assaying transcription factor activity directly rather than relying on binding events for proxy (Cusanovich et al., 2014).

Is BLMP-1 a core oscillator gene? Clearly, oscillations occur even in the absence of BLMP-1, and survival is possible under some conditions (i.e., on plate), if greatly impaired under others (in microwell plates in a luminometer as shown here, or in microfluidic chambers, as reported elsewhere (Stec et al., 2021)). Intuitively, this might be taken to argue against a core oscillator function of BLMP-1. Yet, individual deletion of most mouse circadian core oscillator genes does not abrogate oscillations either and instead alters the oscillation period – indeed typically by a rather modest amount corresponding to < 5% of the wild-type period (Hogenesch and Ueda, 2011). Hence, the finding that BLMP-1 depletion alters the oscillation period would be well consistent with its being a core oscillator component (Takahashi, 2004). Intriguingly, its direct targets include *nhr-23* and *nhr-25*, additional hits from the previous screen (Meeuse et al., 2022) and known molting factors (Gissendanner et al., 2004; Gissendanner and Sluder, 2000; Kostrouchova et al., 1998; Kostrouchova et al., 2001; Kouns et al., 2011). Future research will reveal whether these factors constitute components of a gene regulatory network that can generate autonomous oscillations.

### Blimp1/Prdm1 functions in temporal hair follicle patterning

Mammalian hair follicles undergo regular cycles of expansion (anagen), regression (ketagen) and quiescence (telogen). Recently, (Telerman et al., 2017) showed that mouse *Blimp1/Prdm1* was dynamically expressed in the dermal papilla compartment of the hair follicle, with high levels specifically observed during anagen induction and through advanced anagen. Moreover, dermis-specific *Blimp1* deletion resulted in a delayed onset, and thereby duration, specifically of anagen. Finally, mirroring the cuticular defects seen in *blmp-1* mutant *C. elegans*, *prdm1* mutant mice exhibit skin barrier defects. Given these striking parallels, we propose that *C. elegans* is a suitable model to study general principles of homeostatic skin regeneration.

Molting is frequently defined as a process of exoskeleton replacement in ecdysozoa (Lazetic and Fay, 2017) that is unique to this group of animals (Ewer, 2005), constituting one of their defining features (Valentine and Collins, 2000). Yet, rhythmic renewal of the integument (outer shell) is seen widely also in vertebrates, as illustrated by periodic skin shedding in reptiles and amphibians, feather shedding in birds, or hair shedding in mammals such cats or rabbits, and it may indeed be universal among animals (Stenn and Paus, 2001).

Our results suggest the possibility of a common basis of vertebrate and invertebrate molting processes and specifically imply that homeostatic skin regeneration may involve similar mechanisms across the animal kingdom.

## Acknowledgements

We thank Gert-Jan Hendriks for advice and discussion during the conception of this project, Stephane Thiry, Kirsten Jacobeit and the FMI Functional Genomics Facility for RNA sequencing, Iskra Katic for help in generating transgenic strains, Kathrin Braun for help with the Western Blots, Lukas Burger and Michael Stadler for advice on ChIP-seq analysis, Laurent Gelman for pivotal help with imaging, and Charisios Tsiairis and Benjamin Towbin for a critical reading of the manuscript.

## Author contributions

Y.P.H. performed all transcriptional reporter RT-qPCR time courses. M.W.M.M. and Y.P.H. performed luciferase assays with *blmp-1* RNAi and mutant animals respectively. M.W.M.M. performed the auxin inducible degron luciferase assays. Y.P.H. performed and analyzed the *blmp-1(tm548)* mutant time course using a realignment approach developed by D.G.. Y.P.H. acquired and analyzed single worm imaging data. H.G., M.W.M.M. and Y.P.H. conceived the project and Y.P.H. and H.G. wrote the manuscript.

## Data and materials availability

All sequencing data generated for this study have been deposited in NCBI’s Gene Expression Omnibus (Edgar et al., 2002) and are accessible through GEO Series accession number GSE141514 (*blmp-1(tm548)* RNA-sequencing), GSE163461 (N2 and *blmp-1(tm548)* RNA-sequencing, 2^nd^ replicate). GSE147388 (N2 RNA-sequencing) provides a wild-type time course collected and sequenced in parallel to GSE141514 that was published previously (Meeuse et al., 2020).

Published research reagents from the FMI are shared with the academic community under a Material Transfer Agreement (MTA) having terms and conditions corresponding to those of the UBMTA (Uniform Biological Material Transfer Agreement).

## Methods

### Worm strains

*HW1370: EG6699; xeSi136[F11E6.3p::gfp::h2b::pest::unc-54 3’UTR; unc-119 +] II* (Meeuse et al., 2020)

*HW1939: EG8079, xeSi296[eft-3p::luciferase::gfp::unc-54 3’UTR] II* (Meeuse et al., 2020)

*HW2532: EG8079, xeSi296[eft-3p::luciferase::gfp::unc-54 3’UTR; unc-119 +] II; blmp-1(tm548) (I)* (Meeuse et al., 2020)

*HW2120: blmp-1(xe80[aid::blmp-1]) I; EG8079, xeSi296[eft-3p::luciferase::gfp::unc-54 3’UTR, unc-119(+)] II; EG8080, xeSi376[eft-3p::TIR1::mRuby::unc-54 3’UTR, cb-unc-119(+)] III* (this study)

*HW2033: bus-8(e2885) X* (Partridge et al., 2008)

*HW3028: EG6699, xeSi517[F16B4.4p::gfp::h2b::pest::unc-54 3’, unc-119+] II* (this study)

*HW3076: EG6699, xeSi296[eft-3p::luc::gfp::unc-54 3’UTR, unc-119(+)] II; EG8080,*

*xeSi376[eft-3p::TIR1::mRuby::unc-54 3’UTR, cb-unc-119(+)] III*

### Transgenic CRISPR strains *aid::blmp-1*

Endogenous tagging of *blmp-1* with the auxin inducible degron (aid) was performed by CRISPR/Cas9 using *dpy-10(cn64)* co-conversion (Arribere et al., 2014). The sgRNA sequence: 5’ gccgaagagaacggtgccgg 3’ was cloned into Not1-digested pIK198 (Katic et al., 2015) by Gibson assembly using the hybridized sequence from **5’** AATTGCAAATCTAAATGTTTgccgaagagaacggtgccggGTTTAAGAGCTATGCTGGAA 3’ and **5’** TTCCAGCATAGCTCTTAAACccggcaccgttctcttcggcAAACATTTAGATTTGCAATT **3’**.

The aid sequence was synthesized by IDT (Integrated DNA Technologies) as a gBlocks® Gene Fragments and contained 65 bp homology to *blmp-1* locus, 30 bp downstream of the ATG startcodon.

**5’**ttcgatctcattttaaacaaaacctgtaaaaaatgGGTCAAGGAAGTGGGGATGACGGTGTTCCGatgcctaaagatccagccaaacctccggccaaggcacaagttgtgggatggccaccggtgagatcataccggaagaacgtgatggtttcctgccaaaaatcaagcggtggcccggaggcggcggcgttcgtgaagCCGGCACCGTTCTCTTCGGCTGCTGCGGCAGCTCACTCACCACCTCATTCTCCCCTTTCTGTCGG **3’**.

The injection was performed in wild-type animals which were injected with 10 ng/μl gBlock, 100 ng/μl sgRNA plasmid, 20 ng/μl AF-ZF-827 (Arribere et al., 2014), 50 ng/μl pIK155 and 100 ng/μl pIK208 pIK198, both from (Katic et al., 2015).

### Transgenic reporter strain generation

To generate strain *HW3028: EG6699, xeSi517[F16B4.4p::gfp::H2B::pest::unc-54 3’, unc-119+] II*, the putative *F16B4.4* promoter was amplified from genomic DNA using primers “promoter FW + OH”: gcgtgtcaataatatcactcatctattatcgttaaatgataactgtagt and “promoter RV + OH”: CCATGGCTAAGTCTAGACATgattgaacaaaatcggaatgatg and inserted into *Nhe*1-digested pYPH0.14 backbone using Gibson assembly (Gibson et al., 2009) as previously described (Meeuse et al., 2020). Transgenic animals were obtained by single copy-integration of the transgene into the ttTi5605 locus (MosSCI site) on chromosome II into EG6699 animals, with the published MosSCI protocol (Frøkjær-Jensen et al., 2012).

### Luciferase assays

Embryos were obtained by bleaching gravid adults that express the *xeSi296* transgene *[eft-3p::luc::gfp::unc-54 3’UTR, unc-119(+)] II* obtained by single-copy integration into the *oxTi185* locus on chromosome II. Single embryos were transferred by pipetting into a well of a white, flat-bottom, 384-well plate (Berthold Technologies, 32505) and hatched in 90 µl liquid culture using S-Basal medium (Stiernagle, 2005). For RNAi experiments, the feeding method was used. *E. coli* HT115 bacteria carrying empty plasmids (L4440, mock RNAi) or an RNAi plasmid with an insert targeting *blmp-1* (Fraser et al., 2000; Kamath et al., 2003) were induced with 1 mM IPTG for 1 hour at 37°C. For *blmp-1* mutant luciferase assays, OP50 was used instead of HT115 bacteria. Bacteria were diluted in S-Basal medium (OD_600_ = 0.9), with 100 μM firefly D-luciferin, 100 μg/ml ampicillin and 1 mM IPTG in the case of RNAi. For Auxin Inducible Degradation (AID) experiments, *E. coli* OP50 were diluted in S-Basal medium (OD_600_ = 0.9) and 100 μM Firefly D-Luciferin (p.j.k., 102111). 3-Indoleacetic acid (Auxin, Sigma-Aldrich, I2886) was dissolved in 100% ethanol and diluted 400 times in the culture medium obtaining concentrations as indicated. Vehicle control condition is 0.25% ethanol. Auxin or vehicle control was included in the culture medium at the start of the assay or was pipetted into single wells during the assay at time points indicated. Plates were sealed with Breathe Easier sealing membrane (Diversified Biotech, BERM-2000). Luminescence was measured using a Luminometer (Berthold Technologies, Centro XS3 LB 960) for 0.5 seconds every 10 minutes for 72 hours at 20°C in a temperature-controlled incubator.

Luminescence data was analyzed using an automated algorithm for molt detection in MATLAB, with the option to manually annotate molts in a Graphical User Interface. In short, the hatch was detected by the first data point that exceeds the mean + 5*stdev of the raw luminescence of the first 20 time points and also exceeds the raw luminescence by 3. The molts were detected according to the method previously described (Olmedo et al., 2015) implemented in MATLAB.

## Hoechst 33258 staining

Synchronized L1 worms were obtained by bleaching gravid adults and hatching the released eggs into M9 buffer (Stiernagle, 2005). Larvae were plated on NGM plates (Stiernagle, 2005)seeded with OP50 bacteria and grown up to the L4 stage at 25°C. Worms were then washed 3 times in 10 ml of M9 buffer. After washing, Hoechst 33258 dye was added to 10 ml to a final concentration of 1µg/ml and incubated for 15min on a rotating wheel. Incubation with Hoechst 33258 was then followed by 3 washes in M9 after which worms were concentrated in 1 ml of M9 of which a few µl were mounted on a 2% (w/v) agarose slide with 3 µl of levamisole (10 mM) before a z-stack (z-stack interval 0.7) was acquired using a LSM700 confocal microscope (Axio Imager Z2 (upright microscope) + LSM 700 scanning head, 40x/1.3 oil immersion objective, 6% laser power, 378 ms exposure time, 512x300 pixels). The z-stack was mean projected and grey values were adjusted the same intensity range for all images (0-8000) in Fiji. Staining was quantified manually by assigning worms as being stained if blue signal in the nuclei was obvious.

## BLMP-1 ChIP-seq analysis

We analyzed the ModENCODE dataset (GSE25803) of BLMP-1 ChIP-seq data to count the reads in promoter regions (defined as 1000bp around the ATG start codon, using WS220/ce10 annotations). We downloaded the raw fastq files (see supplementary table 1) and performed the alignment using the function *qAlign(samples, genome = genomeName, pair=’no’, clObj = cl)* from the QuasR package (Gaidatzis et al., 2015) in R to the *C. elegans* genome version ce10 since our RNA sequencing time course was also aligned to ce10 (see above). We used the resulting bam files as input for MACS2 (Zhang et al., 2008) using the command *macs2 callpeak -t ChIP_samples -c control_samples –outdir output_directory -n ChIPvsCtrl_BLMP-1 -g ce -f BAM.* To count the reads within the promoter regions we used the function *qCount(proj3, promoters_of_coding_transcripts, clObj = cl)* from the QuasR package (Gaidatzis et al., 2015) in R and normalized the counts to the mean library size. To calculate the enrichment of counts in promoters between ChIP and control samples we added a pseudocount of 8, log_2_ transformed the counts and calculated the enrichment using *log_2_(promoter_counts_ChIP+8) - log_2_(promoter_counts_control+8).* The mean enrichment of the two replicates per gene was used to threshold the genes of interest at a mean enrichment of 2-fold (log_2_ of 1).

## blmp-1(tm548) mutant time course

Synchronized L1 worms, obtained by bleaching gravid adults and overnight incubation of released eggs in M9 (Stiernagle, 2005), were plated on NGM plates with OP50 bacteria and grown at 25°C. Samples were taken hourly from 1 hour until 24 hours of development and RNA isolation was performed using conventional RNA isolation with phenol chloroform extraction (adapted from (Bethke et al., 2009). Sequencing libraries were prepared using the TruSeq Illumina mRNA-seq (stranded - high input) protocol followed by sequencing using the HiSeq 50 Cycle Single end reads protocol on HiSeq 2500.

### Processing of RNA sequencing results

RNA-seq data were mapped to the *C. elegans* ce10 genome using STAR (Dobin et al., 2013) with default parameters (version 2.7.0f) and reads were counted using htseq-count (Anders et al., 2015) (version = 0.11.2).

Counts were scaled by total mapped library size for each sample. A pseudocount of 8 was added and counts were log_2_-transformed and then quantile normalized using *normalize.quantiles()* from the *preprocessCore* (version 1.52.1) (Bolstad, 2021) in R.

### Realignment of blmp-1 mutant RNA sequencing time courses

In order to compare equivalent developmental time points, we had to realign the wild-type to the *blmp-1* mutant time course samples to account for differences in developmental tempo (Fig. S3). We realigned the time courses using the log_2_-transformed gene expression of all oscillating genes (n = 3’739). The mean normalized, 10 times oversampled and zero padded time series of each gene of the faster time course(s) were interpolated using a Lanczos kernel (a=2):

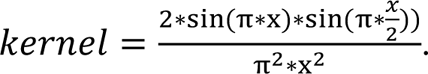

We then calculated the errors of the interpolated time points with the non-interpolated data of the slowest time course in log_2_ (Fig S3E), in our case the *blmp-1(tm548)* mutant samples of replicate 2. For each time point of the slowest time course, we selected the interpolated gene expression values of the time point with the smallest sum of the squared errors in both wild-type replicates and the *blmp-1(tm548)* replicate 1 to create a new realigned time course with the same number of time points as in the slowest condition (24 time points). Since the rhythmic process of oscillations leads to repetitive small errors, we had to limit the scanning for smallest error to the diagonal (Fig S3E).

#### edgeR using realigned data

In order to identify differentially expressed genes, we used edgeR (McCarthy et al., 2012; Robinson et al., 2010) on the realigned data. We first created counts from the log_2_-transformed realigned data again and ensured only non-negative counts by using pmax(2^(realigned_log_2__data) -8,0) together with the real count data from the *blmp-1(tm548)* replicate 2. We kept expressed genes by selecting genes which had at least 3 time points with counts higher than 5. These count data served as an input for edgeR (version 3.32.1). We then used the design matrix model.matrix(∼time + batch + time:treat) to estimate the dispersion, run a QL fit and performed an F-test using *glmQLFTest(fit, coef = coefficients_for_time_batch_time:treat)*. Since we tested many time points, the obtained p-values were generally small and we would be able to obtain even small changes between samples by choosing a conventional p-value cutoff of 0.05. Based on this circumstance, we selected genes not only based on a p-value but additionally also based on the mean of the absolute log-fold changes per gene to identify significant differentially expressed genes (Fig S4A), resulting in our differentially expressed list of 1’655 genes (Fig 2A). The p-value threshold for identifying genes was set at: *-log10(p-value) = 25*8^(-mean(abs(fold_changes) + 5*.

We ordered the gene expression and the fold changes of these 1’655 genes according to a 1-dimensional tSNE performed on the fold changes of both replicates using the function *Rtsne(fold_changes,dim=1,pca=FALSE,perplexity=100,theta=0.25)* from the Rtsne package (Krijthe, 2015) in R and used the first tSNE from the output for ordering (Fig 2A).

#### Amplitude and period calculation using Hilbert transform

Calculation of amplitudes and periods was performed as described in (Meeuse et al., 2020) in Python. Published wild-type peak phases for annotation were from (Meeuse et al., 2020).

### Single worm imaging

Sample preparation and analysis were performed as described in (Meeuse et al., 2020).

For scaling the GFP intensities in Fig 2 and S4 we used python with the numpy package (version 1.16.4). For Fig 2D, we scaled all single GFP traces from hatch until molt 4 exit to 200 datapoints (=100%) using the function *interp()* from the numpy package (version 1.16.4). We then plotted the scaled values using seaborn’s function *lineplot()*, using the standard deviation as error (*ci = “sd”*). In Fig S4F,G we scaled all individual GFP intensity traces to the absolute mean duration of mock and *blmp-1* RNAi separately for each larval stage in hours.

### Supplementary Figures

**Fig S1:**
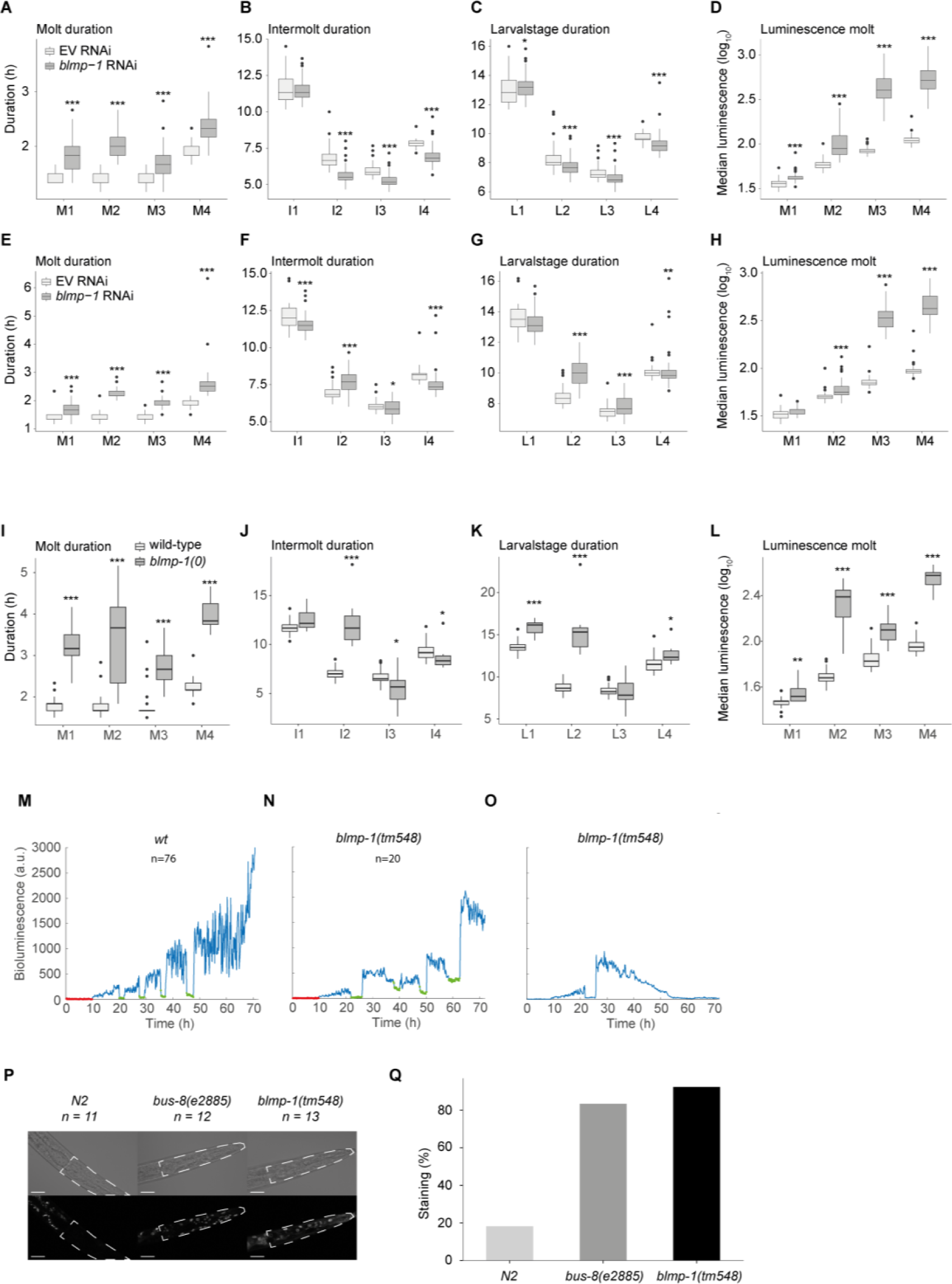
Replicates of experiments revealing developmental phenotypes upon BLMP-1 loss, Related to Figure 1. (**A-C, E-G, I-J**) Boxplots of the quantification of single animal molt, intermolt and larval stage durations in mock and *blmp-1* RNAi or wild-type and *blmp-1(0)* mutant animals as indicated. (**D,H,L**) Boxplots showing log10-transformed luminescence intensities during the indicated molts in mock (EV) and *blmp-1* RNAi (D, H) and in wild-type and *blmp-1(tm548)* mutant animals (L). Both replicates confirm the longer molts (**A, E**), that are accompanied by shorter intermolts (**B**), or similar intermolt lengths (**E, F**), leading to similar (**C**) or slightly longer (**G**) larval stage durations in *blmp-1* RNAi compared to mock RNAi. Luminescence values are increased in both replicates during the molts (**D, H**). Boxplots represent the median as horizontal line and hinges extend to first and third quartiles, the whiskers extend up to 1.5*IQR (interquartile range). Data beyond that limit is plotted as dots. Significantly different durations are indicated (* P<0.05, ** P<0.01, *** P<0.001, Wilcoxon unpaired, two-sample test). **M – O,** Representative luminescence traces from a wild-type (M, n=80) and surviving *blmp-1(tm548)* mutant (N, n=7). Many *blmp-1(tm548)* mutant animals exhibit a gradual decrease of luminescence after the exit from the first molt (O), presumably reflecting death. **P, Q,** Animals of the indicated genotypes were incubated with Hoechst 33258 to test for cuticle permeability Representative images of (P) and quantification of the percentage of stained worms (Q) are shown. A dashed white line in (P) outlines the head region used for scoring; note that signal outside this area in N2 corresponds to autofluorescence. Scale bars in (P) represent 20 µm, n≥11 in (Q).

**Figure S2:**
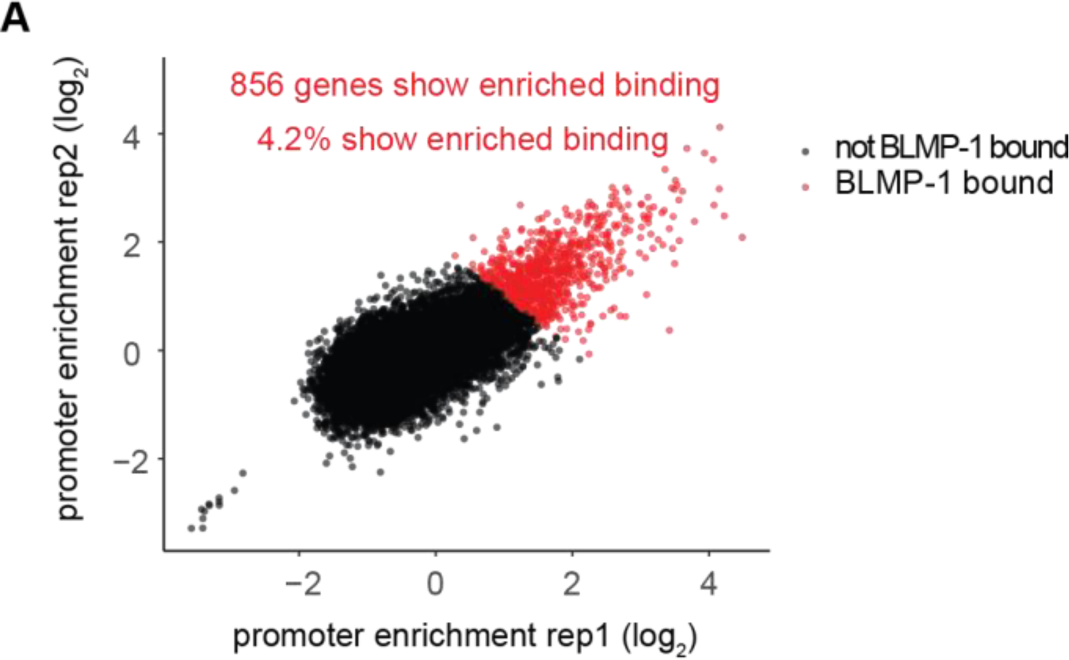
BLMP-1 binding is enriched on oscillating genes, Related to Figures 2 – 4. **A**, Log_2_-fold enrichment of BLMP-1 binding to promoters (1000bp around the ATG start codon) for all coding genes (n = 20,392). Identified genes that show enriched binding (on average log_2_ > 1 across the replicates) are labelled in red (n = 856).

**Fig S3:**
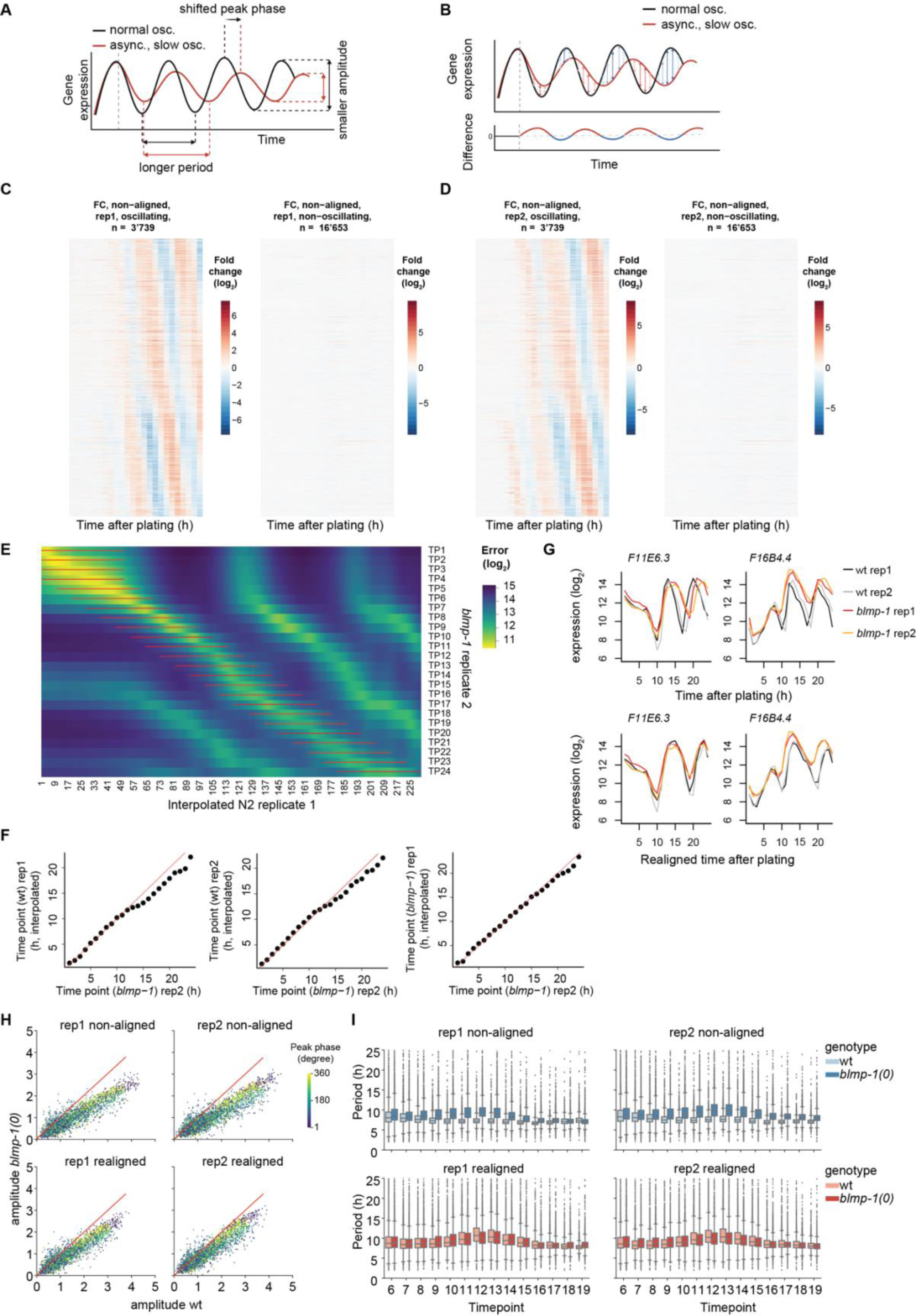
Temporal realignment corrects for different developmental tempos among strains, Related to Figure 2. **A**, Model of changes in oscillatory gene expression upon slower and unsynchronous development of a mutant (red) in comparison to a wild-type strain (black). In *blmp-1* conditions, genes still coupled to development will slow down together with slower development. Furthermore, asynchronous development will result in reduced amplitudes in *blmp-1* conditions. Created with Biorender.com **B**, Resulting hypothetical rhythmic fold changes from a situation as in A. **C, D**, Measured mean-centered fold changes of oscillating (n = 3’739) and non-oscillating (n = 16’653) genes in log_2_ in two RNA sequencing experiments, comparing *blmp-1(tm548)* mutants with wild-type. Notice lack of overt differences in expression before 10 h of development and wide-spread and rhythmic fold changes in expression that become apparent thereafter, consistent with the altered developmental rates observed in Figures 1 & S1. **E**, After spline interpolation of the time courses, the sum of the squared error between the interpolated time course(s) and the *blmp-1(tm548)* mutant replicate 2 original log_2_-transformed gene expression profile is calculated for all oscillating genes. For every time point of the *blmp-1(tm548)* replicate 2, the corresponding interpolated time point is chosen by identifying the smallest error. Since repetitive small errors occur due to the nature of the underlying oscillating gene expression, we limit the search to a diagonal sub-selection, indicated in red. **F**, Calculation of the interpolated time point (h) that corresponds best to the *blmp-1(tm548)* replicate 2 timepoints (h). **G**, log_2_-transformed gene expression of *F11E6.3* and *F16B4.4* before (top) and after (bottom) temporal realignment of the gene expression profiles of two replicates each of N2 wild type (black and gray traces) and *blmp-1(tm548)* mutant animals (orange). **H**, Mean amplitudes calculated from the Hilbert transform of Butterworth-filtered signals (as in Meeuse et al., 2020) from time points 10 - 20 (to circumvent edge effects) of non-aligned and realigned log_2_-transformed expression data of oscillating genes. Color indicates the peak phase of the individual genes plotted. **I**, Period over time calculated from the Hilbert transform of Butterworth-filtered signals (as in Meeuse et al., 2020) from time points 7 to 20 to circumvent edge effects.

**Figure S4:**
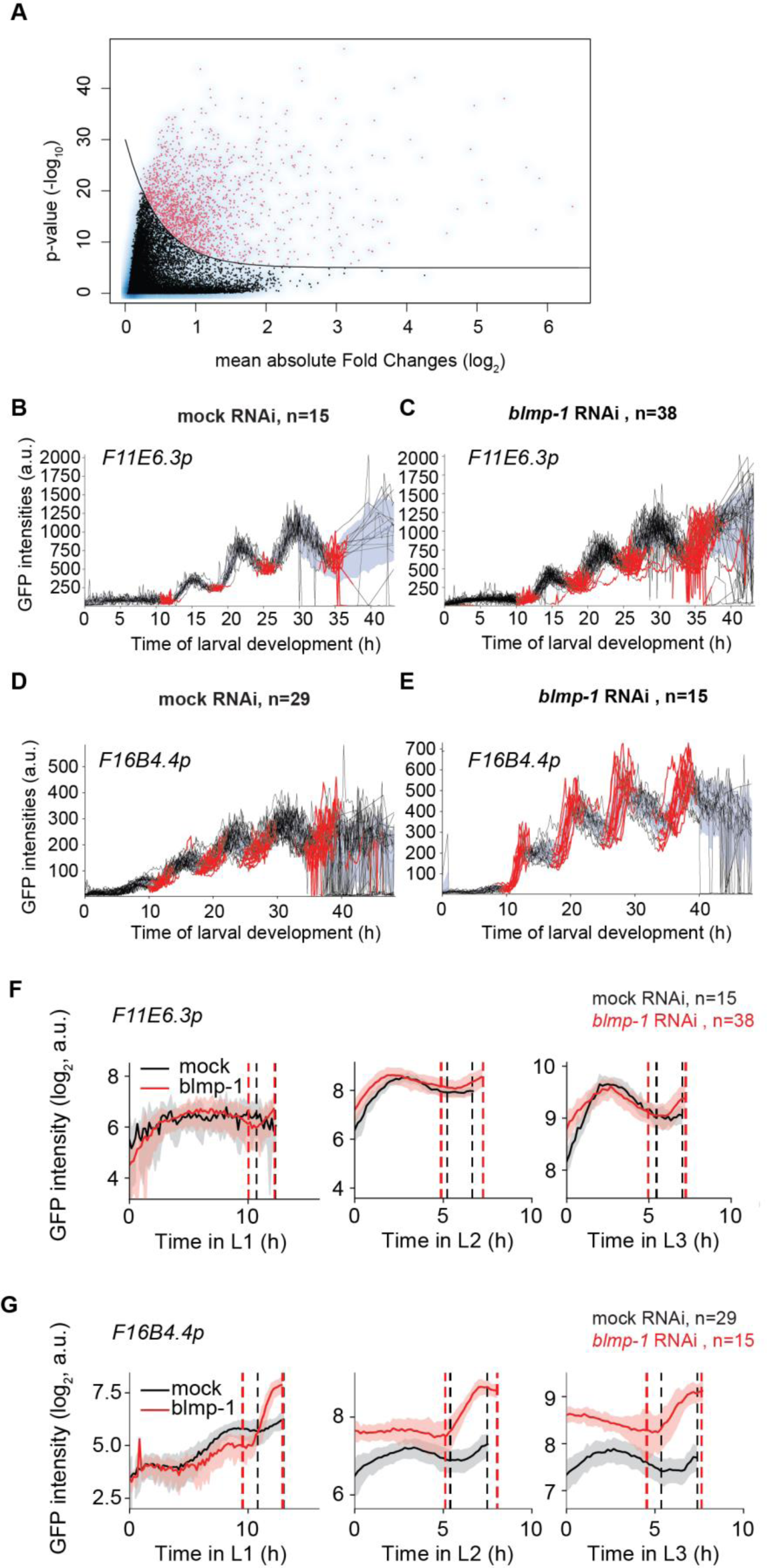
Identification of differentially expressed genes in *blmp-1(0)* mutant animals, Related toFigure 2. **A,** Scatter plot of -log10 p-values over mean of absolute fold changes between *blmp-1(tm548)* and N2 wild type samples over the course of development using the temporally realigned gene expression data from Figure S4. Dots represent individual genes, with genes above the indicated threshold line (Methods) colored in red. **B-E,** Unscaled GFP intensities of single worm image analysis of *F11E6.3p::gfp* (B), (C) and *F16B4.4p::gfp* (D), (E) reporters in mock RNAs (B, D) or *blmp-1(RNAi)* treated animals. t = 0 h corresponds to hatch. Molting times are indicated in red.For missing time points we interpolated linearly, mostly after the fourth molt. **F, G,** Per-larval-stage scaled log_2_-transformed mean GFP intensities of single worm image analysis from B-E. The GFP intensities were scaled to the mean larval stage of mock and *blmp-1* RNAi separately for L1-L3 for the *F11E6.3p::gfp* (F) and the *F16B4.4p::gfp* (G) reporters. Vertical dashed lines indicate beginning and end of molt. The standard deviation is shown in shading.

**Fig S5:**
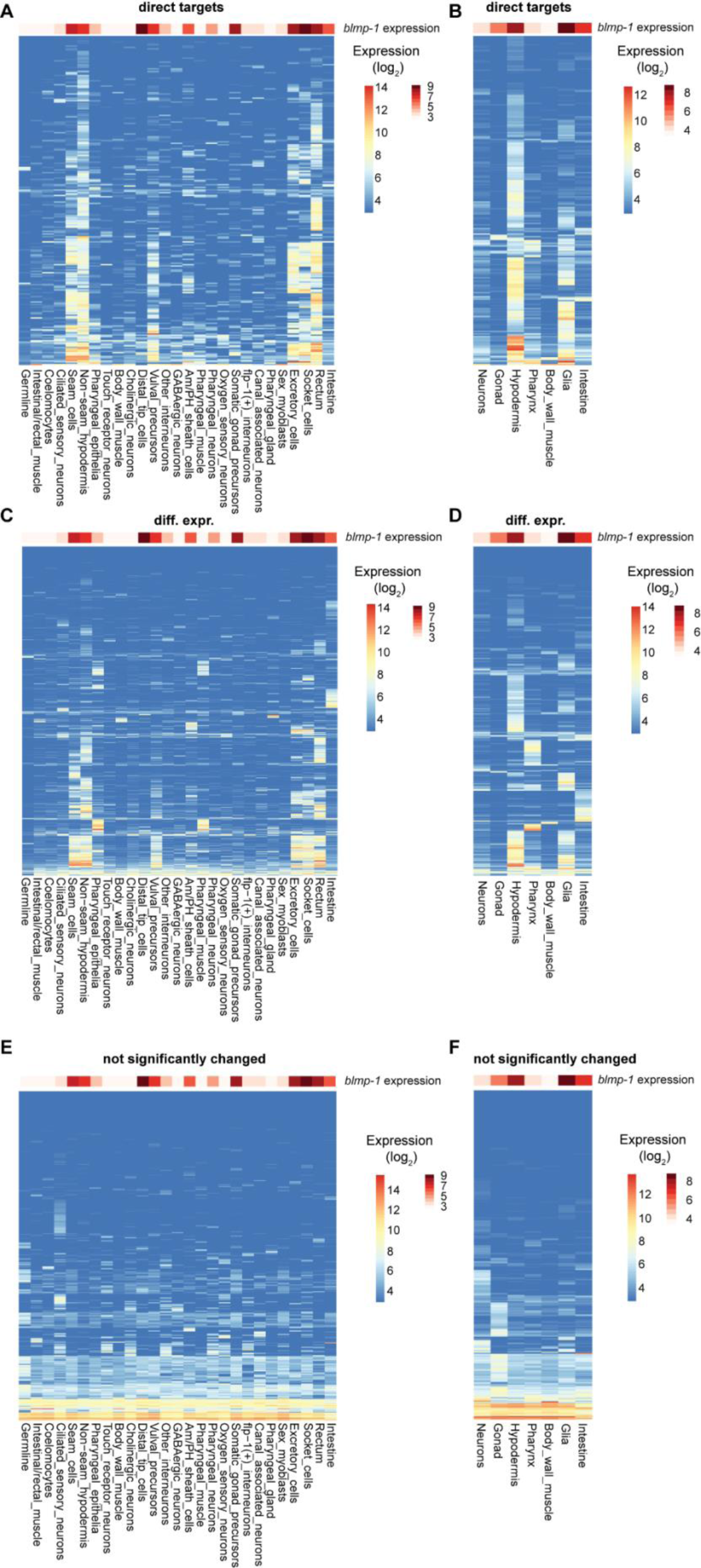
Spatial expression patterns of different gene classes, Related to Figure 3. Heatmaps are generated using single-cell RNA sequencing data from (Cao et al., 2017) **A**, **B**, Expression of direct BLMP-1 target genes at the cellular (A) and tissue (B) level. **C**, **D,** As in (A), (B) but for genes that are differentially expressed in *blmp-1* mutant versus wild-type animals. **E, F**, As in (A), (B) but for genes that are not differentially expressed in *blmp-1* mutant versus wild-type animals.

**Fig S6:**
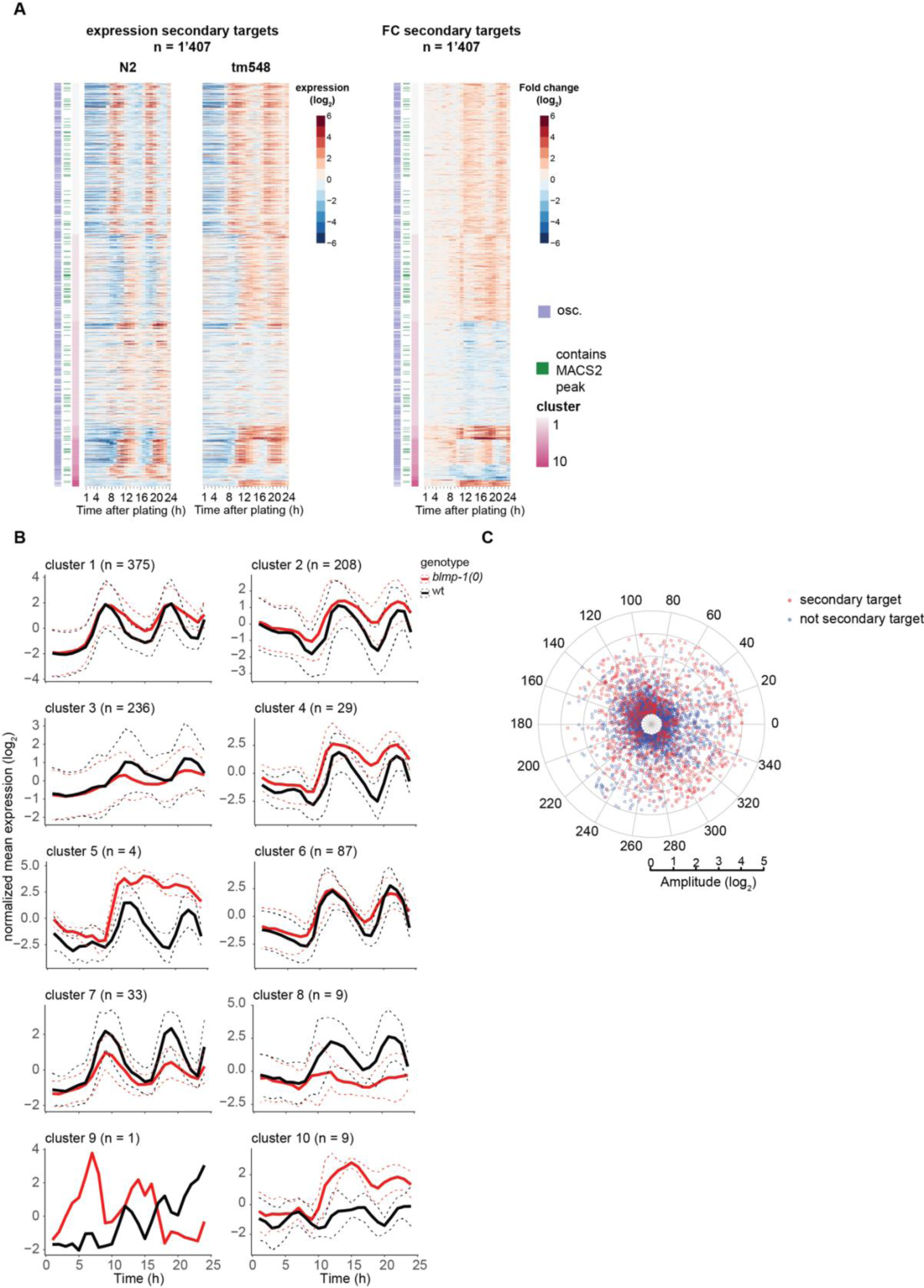
Secondary targets are affected in oscillatory gene expression but do not display phase specificity. Related to Figure 4. **A,** Heatmap of log_2_-transformed gene expression and fold changes of secondary BLMP-1 targets. Columns on the left indicate oscillating genes (violet), genes for which MACS2 calles a BLMP-1 binding peak in the ChIP-seq data (green), and expression clusters (shades of purple). **B**, Mean-normalized mean expression and peak phase distributions of oscillating secondary target genes according to the 10 clusters in A. The dashed lines represent the standard deviation. **C**, Peak phases (circular axis, degree) and amplitudes (radial axis, log_2_) of secondary target genes (red) and the remaining oscillating genes (blue).

